# Ventricular Forebrain Organoids Reproduce Macroscale Geometry of the Developing Telencephalon

**DOI:** 10.64898/2026.03.17.712213

**Authors:** Alexander W. Justin, Alexander Anderson, Luca Guglielmi, Madeline A. Lancaster

## Abstract

During development, the size of the neuroepithelial cell pool plays a key role in establishing brain size, determining the numbers of derived progenitors and subsequent neuronal cell types. While early histogenesis is well modelled in brain organoids, the organ-scale geometry of the telencephalon is not accurately recapitulated. Herein, we present a new approach for generating ventral and dorsal forebrain organoids which develop a large ventricular neuroepithelium, characteristic of the closed telencephalic vesicle. Using a growth medium that supports aerobic glycolysis and is typically used for endothelial cells, we modulate neuroepithelial expansion to induce a more anatomically accurate neuroepithelial layer which, upon maturation, thickens physiologically to generate the typical neurogenic layered architecture. In addition, we present a new method for embedding organoids in miniature collagen spheres which mimics native extracellular matrix, stabilizes the ventricular geometry for dynamic culture conditions, and provides a means for incorporating vascular cells for neurovascular development. Finally, we demonstrate that human organoids grown under these conditions exhibit dramatically enlarged ventricles and delayed maturation compared to mouse. Together, this approach provides a model of the forebrain neuroepithelium with morphogenetic macroscale geometry and tissue architecture, suitable for investigating neurodevelopment and disease.

## 1. INTRODUCTION

The developing brain forms an intricate architecture around a large fluid-filled cavity. The most anterior part of this neural tube, the prosencephalon, develops by rapid expansion of the neuroepithelium which subsequently undergoes thickening and distinct regional and layered patterning of progenitor and neuronal cell types^1^. Simultaneously, neural crest- and mesoderm-derived precursor cells form a pial surface around the neural tube, through which vascular cells invade and form a nascent blood vessel system following a highly programmatic patterning process. This initial period of neuroepithelial expansion is crucial in establishing mature brain size as it determines the founder progenitor pool available for neurogenesis^2,3^. By modulating neuroepithelial expansion, one can highlight evolutionary differences in brain size and further understanding of cerebral diseases, notably microcephaly and lissencephaly, in which there is an early perturbation to the tissue architecture and organ geometry.

Current brain organoids do not recapitulate the organ-scale geometry of the developing anterior neural tube, yielding highly variable, anatomically inaccurate structures and many miniature rosettes within a single organoid^4^. These small buds are challenging to distinguish^5^ and prevent proper expansion, with lumen size reported to relate to cell shape and cell fate^6,7^. There has recently been interest in generating neural tube architectures using bioengineering approaches such as microfluidics and micropatterning^8,9^. While the presence of a ventricular lumen is visible using these approaches, the enlarged geometry of the prosencephalic and, later, telencephalic vesicles seen in vivo remains to be demonstrated. In addition, producing a telencephalic organoid with clearly defined, invasive vasculature remains a key challenge in the field. For the forebrain, vessels first invade through the pial surface of the ventral forebrain and spread tangentially to reach the dorsal forebrain, forming a subventricular vascular plexus (SVP)^10^. This vascular ingress closely coincides spatiotemporally with neural precursor differentiation as vasculature greatly increases the flow of oxygen and metabolites into the brain, supporting the generation of highly oxygen consumptive neuronal cell types^11^. Recapitulating this neurovascular architecture in 3D cell assemblies would not only enable more complex and mature neural circuits to be produced but also provide a key tool in understanding neurovascular diseases and potential treatments.

Herein, we present methods for generating a neuroepithelial tissue model which shows an enlarged ventricular telencephalon, similar to the telencephalic vesicle *in vivo*. By using a low serum medium with an atypical basal composition usually used for growing endothelial cells (EGM), we produce a large ventricular forebrain organoid from mouse pluripotent cells. By embedding such organoids in miniature collagen hydrogel spheres, mimicking the surrounding extracellular matrix (ECM) of the neural tube while supporting high metabolite exchange, we show that upon removing EGM, the ventricular organoids differentiate to form a thickened telencephalon, consisting of stratified layers, while preserving the ventricular geometry. The establishment of more anatomically correct morphogenesis in organoids enables us to further introduce vascular cells over the large basal surface, mimicking the overlying pial vasculature *in vivo*. However, difficulty in achieving vascular ingress into the telencephalic wall highlights a resistance to infiltration even at such an early stage of brain development. Finally, we demonstrate the application of this method to human cells, revealing species-specific expansion and delay resulting in human organoids with much larger ventricles than mouse, reminiscent of in vivo differences.

## 2. RESULTS

### 2.1. EGM culture reproducibly generates large ventricular structures in mouse neural organoids

Cerebral organoids typically develop many small protruding neuroepithelial buds or rosettes but lack large continuous ventricles at early stages. We included an additional culture stage using EGM with Growth-Factor Reduced Matrigel (MG) between neural induction and neuroepithelial differentiation, which enabled formation of a continuous neuroepithelium in mouse forebrain organoids with a dorsal or ventral identity (Fig. 1a, b). Over the course of several days, the outer layer of the organoid detached from the central mass forming a thin, well-defined, and polarized neuroepithelium, reminiscent of the early anterior neuroectoderm (Fig. 1c). By Day 7, large ventricular spaces were visible and the neuroepithelium attached only sporadically to the central mass. When compared to standard culture protocols using Maturation medium (MAT) from the post-neural induction stage, EGM-derived mouse organoids displayed significantly fewer but larger neuroepithelial buds (Fig. 1d, e). Measuring individual buds, EGM organoids exhibited significantly larger visible ventricular lumen areas, distinct from the central organoid mass, with significantly longer neuroepithelial surfaces when measured in cross-section (Fig. 1f – left, centre). However, the average thickness of the neuroepithelium remained unchanged compared to MAT-derived organoids (Fig. 1f – right).

**Fig 1.**
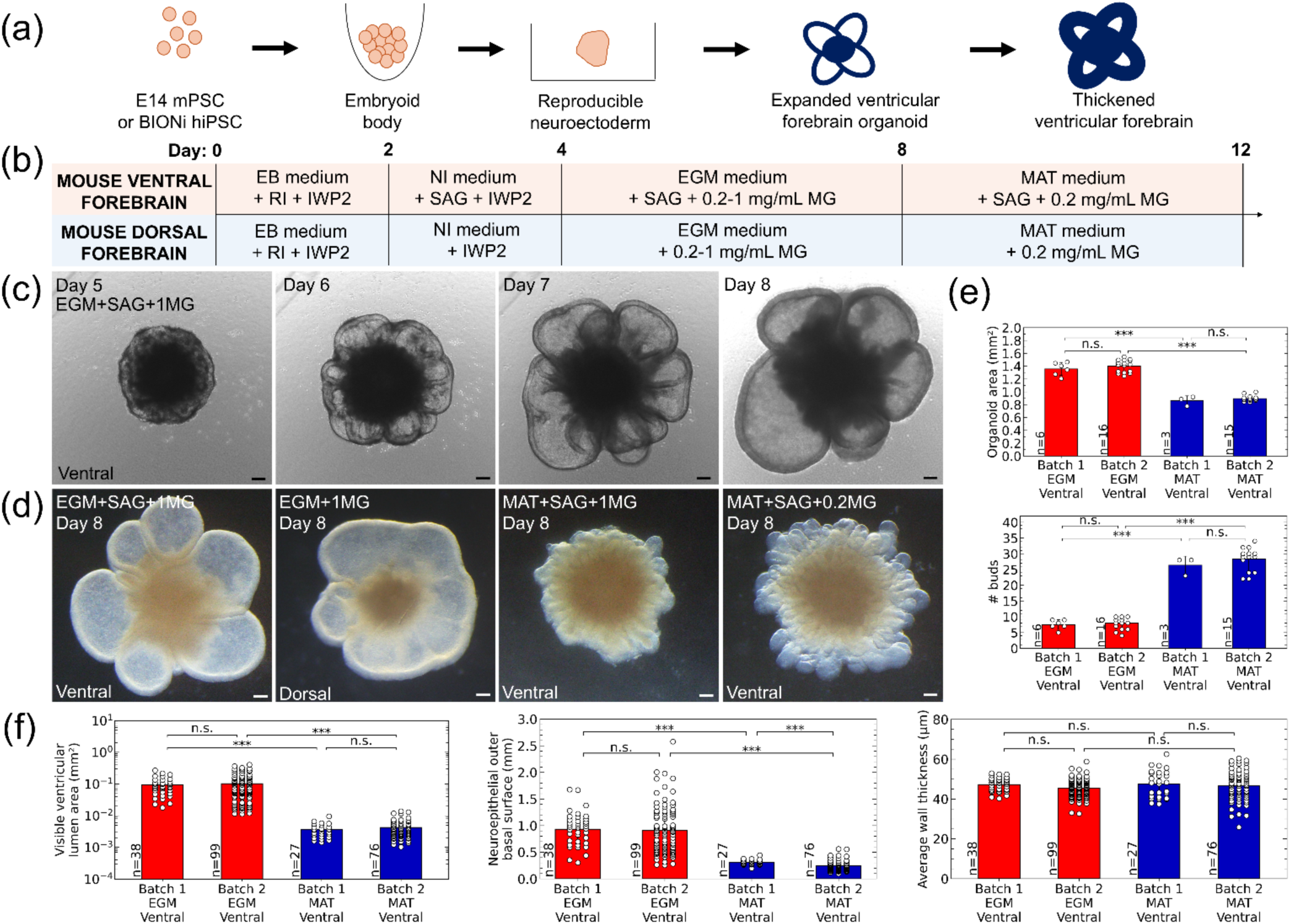
EGM culture reproducibly generates ventricular tissue architecture in mouse forebrain organoids. **(a)** Summary schematic for generating well-defined forebrain ventricles using EGM. **(b)** Protocol for generating ventral (guided) and dorsal (unguided) mouse forebrain organoids. **(c)** Following neural induction, mouse neuroepithelial cells undergo proliferation and expansion into a shell-like structure away from the organoid core. **(d)** Mouse ventral and dorsal forebrain organoids display fewer larger buds when treated with EGM as compared those cultured with standard MAT medium, which present miniature and more numerous ventricles, within a contiguous central mass. **(e)** Measurements of organoid area and bud number show significant differences between EGM and MAT culture conditions. **(f)** Measurements of individual buds show significant differences between EGM and MAT culture conditions for visible ventricular lumen cross-sectional area (left) and length of outer basal surface in cross-section (middle), while displaying no significant difference in average neuroepithelial wall thickness (right). EB: embryoid body; NI: neural induction; RI: ROCK inhibitor; IWP2: Wnt production inhibitor-2; EGM: endothelial growth medium; MAT: maturation medium; 1MG: 1 mg/mL growth factor reduced (GFR) Matrigel (MG). Statistics: Levene’s test followed by Kruskal–Wallis with Games–Howell post-hoc comparisons. p < 0.05 (*), p < 0.01 (**) and p < 0.001 (***); n.s. means no statistically significant difference between groups was detected.

### 2.2. EGM culture leads to neuroepithelial expansion without differentiation

To elucidate the process by which neuroepithelial expansion occurs in EGM medium, we tested two other endothelial cell basal media: MCDB131 and Medium 200. While the components of the basal medium of EGM are proprietary, the basal components of the other media are known. By adding the supplements of EGM to the other cell media, we were able to induce the same characteristic neuroepithelial expansion (Fig. 2a), suggesting similar basal medium components. To test whether the supplemental mitogenic growth factors (EGF, IGF, bFGF) were responsible for the observed neuroepithelial bud expansion, we compared EGM culture with and without supplementation, alongside the fully defined Improved Differentiation Medium (without vitamin A, IDM-A) usually used when generating cerebral organoids, with and without the same supplements. Despite the EGM organoids being notably smaller in size, we found the characteristic enlarged buds were present when using EGM basal medium even without supplements (Supplementary Fig. S1a). Instead, IDM-A with the supplements usually added to EGM produced enlarged organoids but without the characteristic neuroepithelial bud expansion (Supplementary Fig. S1b). Finally, the addition of the mitogen bFGF to IDM-A at 1x and 5x concentrations found in EGM did improve bud growth as expected from the literature^12,13^, though the large bud expansion of EGM culture was not detected (Supplementary Fig. S1c).

**Fig 2.**
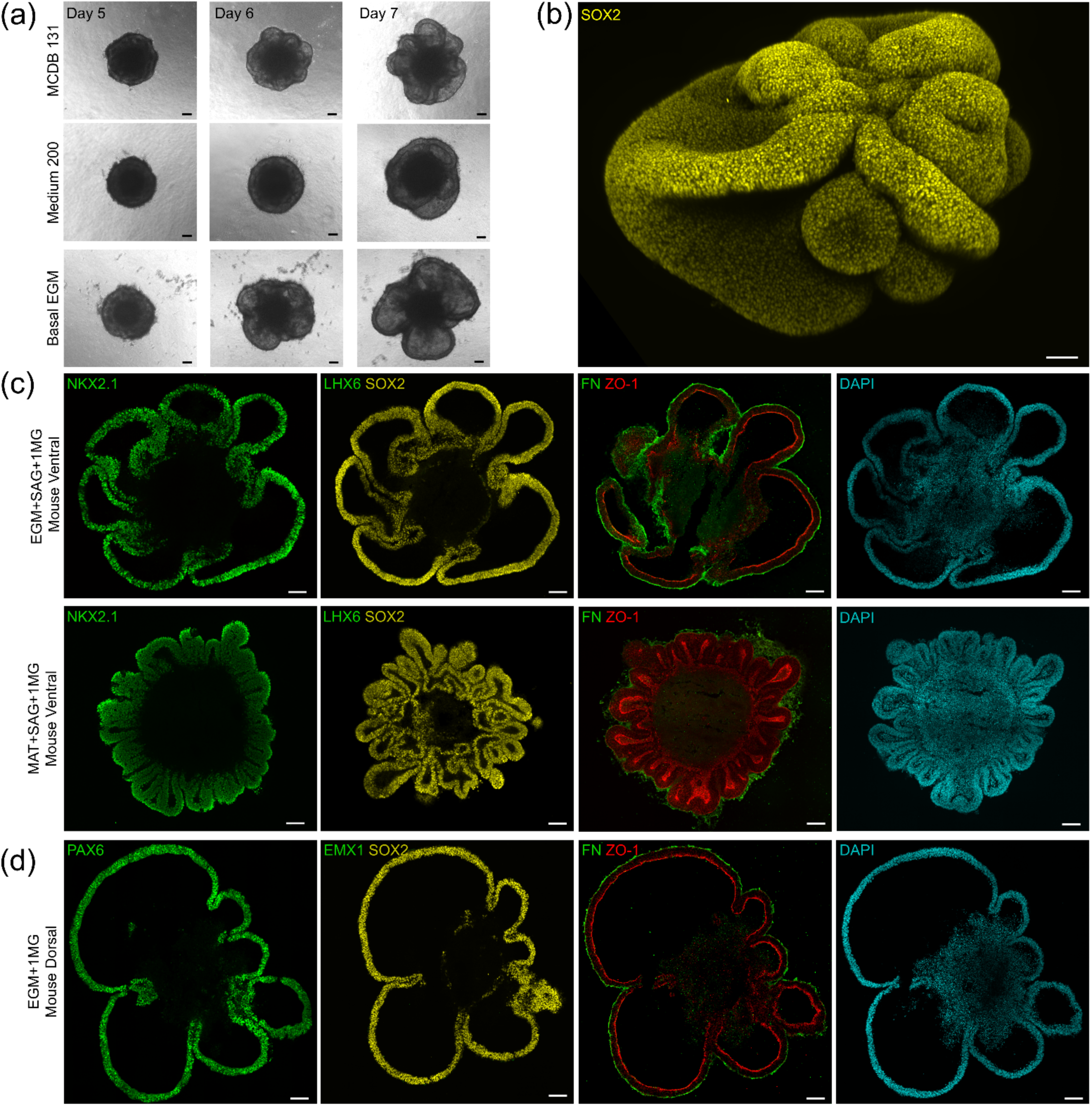
Characterisation of EGM-expanded mouse forebrain organoids. **(a)** EGM supplements were added to common vascular cell basal mediums -MCDB131, Medium 200, and basal EGM -which subsequently exhibit characteristic neuroepithelial layer expansion. **(b)** Whole mount staining of ventral forebrain organoid demonstrates undifferentiated neuroepithelial progenitors marked by SOX2. **(c)** Ventral forebrain organoids with EGM exhibit similar cell identities to MAT-cultured organoids from Day 4-8. However, ventricular lumen and apical-basal surfaces (ZO-1-FN) are significantly better defined within EGM organoids. **(d)** Dorsal forebrain organoids also demonstrate large bud formation away from central organoid mass. DAPI stain reveals pyknotic nuclei characteristic of necrotic core. Scalebars: 100 µm.

The various endothelial culture media have similar nutrient compositions, focused on supporting aerobic glycolysis, which endothelial cells are known to rely on, even in the presence of oxygen, suggesting the metabolic effects may be responsible. Several components of endothelial basal mediums are also known to cause a metal-induced hypoxia-like response, and to stabilize HIF-1α as its master regulator (Supplementary Table 1). To test whether this may be responsible for the expansion, we performed immunostaining for HIF-1α but found a lack of nuclear-localised (active) HIF-1α at Day 8 in EGM media (Supplementary Fig. S2a). We also found that organoids below a certain size threshold of about 1000 cells per organoid – below which would have no discernible necrotic core – did not exhibit distinct neuroepithelial bud expansion (Supplementary Fig. S2b). Necrosis, as a unprogrammed form of cell death, can be driven by a lack of oxygen and nutrient delivery to the core of organoids^14^. We hypothesized that the necrotic core provides a non-adhesive substrate to living cells at the organoid periphery, promoting a hollow tissue architecture and supporting neuroepithelial monolayer formation. However, we did not observe complete detachment of the outer shell neuroepithelium from the core; the enlarged buds remained sporadically attached to the peri-necrotic region within the organoid (Supplementary Fig. S2c). We also showed that EGM-mediated neuroepithelial expansion does not occur by incubation in hypoxic 1% or 2.5% oxygen tensions for 24h (Supplementary Fig. S3). Thus, we concluded that neuroepithelial expansion more likely occurs due to the metabolic effects of the medium components favouring glycolytic metabolism, rather than hypoxia.

Whole-mount (Fig. 2b) and serial sectioning of EGM-cultured ventral mouse organoids (Fig. 2c) revealed that the neuroepithelium remained in an immature state, preserving a ventral forebrain (NKX2.1) identity while showing no signs of neurogenesis by Day 8 (SOX2+, LHX6-). A large and connected neuroepithelial surface was observed in the EGM organoids, while the MAT-derived organoids showed smaller and more numerous buds. Similar markers were observed for dorsal mouse organoids at Day 8 (Fig. 2d), maintaining a dorsal forebrain (PAX6) identity without transition yet to neurogenic pallium (SOX2+, EMX1-), along with smooth, continuous basal and apical surfaces (FN1 and ZO-1). The basal and apical markers also outlined a large, clear ventricle separated from the necrotic core (pyknotic nuclei detected via DAPI stain) which did not occur in MAT-cultured organoids.

We also investigated an assembloid approach using EGM-cultured forebrain organoids to generate a dorsoventral axis within a single neuroepithelial bud (Supplementary Fig. S4). However, we observed that the dorsal and ventral organoids did not fuse sufficiently well to form singular buds and once the medium was switched from EGM to MAT, the differentiating assembloid was far more disorganised in structure than the separate dorsal or ventral organoid equivalents.

### 2.3. Tissue architecture is preserved upon differentiation using expanded ventricular forebrain organoids

Subsequent differentiation of EGM-expanded mouse organoids was investigated first under static culture conditions (Fig. 3a, Supplementary Fig. S5a). A continuance of EGM culture maintained a thin-walled neuroepithelium, but without a continued spatial expansion as observed from day 4 to 8. When compared to organoids cultured in MAT medium from Day 4 onwards, those initially cultured in EGM developed as large singular ventricles, with well-defined apicobasal wall thickening. The ventricles were defined by a fluid-filled space within the organoid and the sharp demarcation of staining for the apical marker ZO-1 (Fig. 3b). Progression to MAT medium supplemented with 0.2 mg/mL MG induced a thickening of the neuroepithelium (Fig. 3a, c). In the case of EGM + 1 mg/mL MG (1EGM), it was necessary to remove the MG that had attached to the organoid to prevent early axonal outgrowth and enable the telencephalic wall to thicken (using Cell Recovery Solution). Immunostaining for differentiation markers displayed clear stratified layers within EGM organoid buds. However, under these static culture conditions, limited disconnected regions of cells were positive for LHX6, a marker of GABAergic neurons originating from the medial ganglionic eminence. While overall geometry and tissue structure was enhanced with this protocol, we did not detect any noticeable differences in the cell type determination between previous EGM and MAT-cultured organoids (Fig. 3d). In addition, the stratified layering of SOX2+ and MAP2+ cells further indicated post-mitotic neurons correctly positioning themselves in the outer layer. When EGM-culture was sustained for Days 8-12, SOX2+ cells were detected through the full thickness of the neuroepithelium, suggesting that EGM could maintain neuroepithelial stemness (Supplementary Fig. S5b).

**Fig 3.**
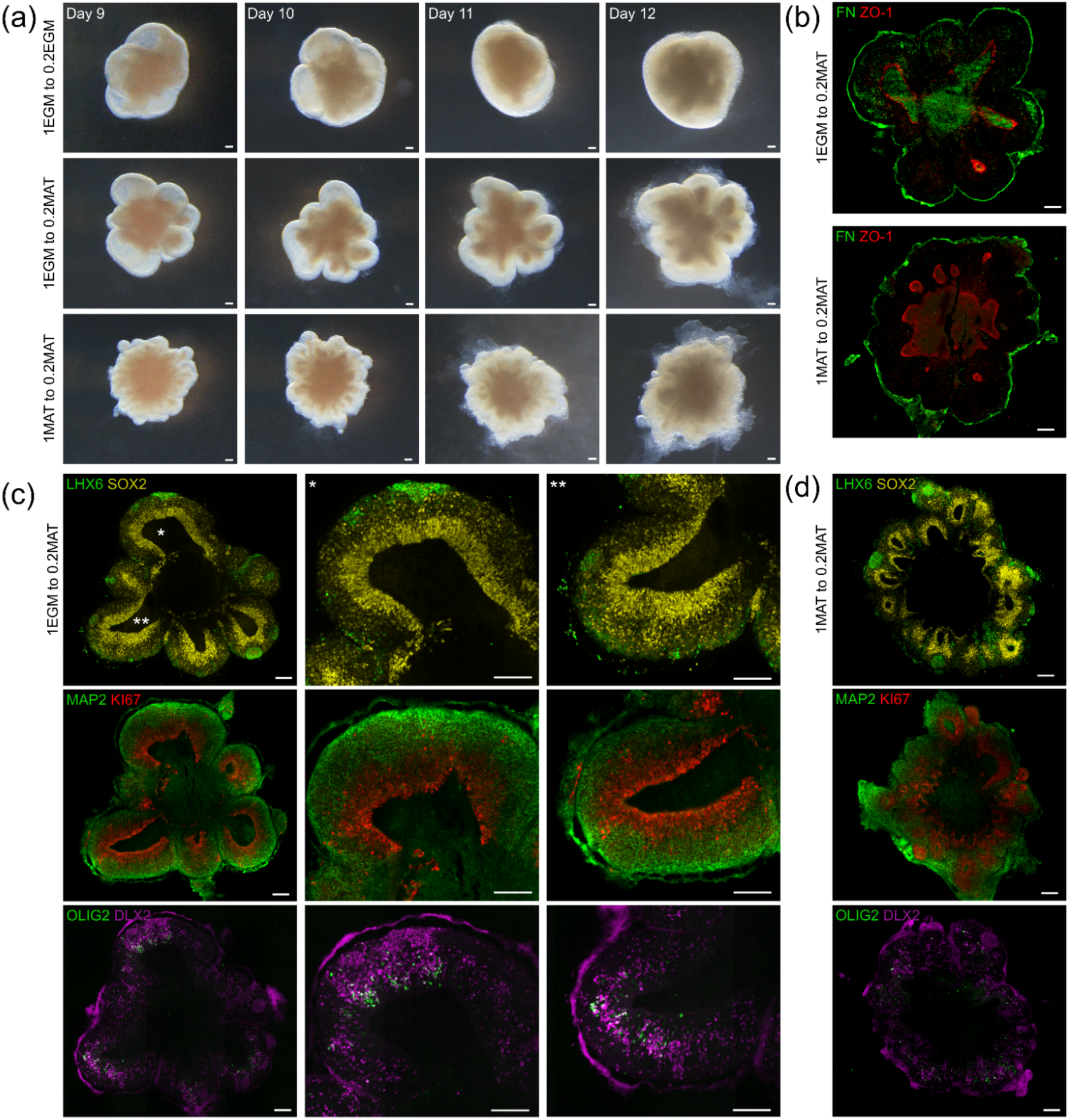
Tissue architecture is preserved upon differentiation using expanded ventricular forebrain organoids. **(a)** Comparison of mouse expanded ventricular organoids statically cultured in EGM or MAT from Day 4-8 in 1 mg/mL GFR MG. Organoids were subsequently statically cultured in EGM or MAT medium supplemented with 0.2 mg/mL MG from Day 8-12. **(b)** FN and ZO-1 staining of Day 12 organoids during differentiation stage maintain well-defined apical and basal surfaces. **(c)** Differentiation markers for EGM-cultured organoids at Day 12, showing large, open ventricular buds and distinct cell layers. Limited differentiation to MGE-derived interneurons, as indicated by LHX6 staining. MAP2 and KI67 staining distinguishes postmitotic neurons from proliferative progenitors. Distinct DLX2 and OLIG2 subpopulations distinguish GABAergic interneuron precursors from multipotent MGE progenitors, mirroring in vivo spatiotemporal development. * and ** provide location of magnified buds (centre and right images). **(d)** MAT-cultured organoids show similar cell populations, though smaller ventricles. Scalebars: 100 µm.

### 2.4. Water-in-oil sphere technique enables small-volume matrix embedding while preserving structure under shaker culture

Dynamic shaker culture provides significantly improved metabolite transfer when compared to diffusion-based, static culture conditions^15^. However, we observed that placing EGM-cultured organoids into shaker culture at Day 8, by which point the neuroepithelium had expanded away from the central organoid mass, led to amorphous growth of the neuroepithelium and a loss of tissue structure (Fig. 4a, Supplementary Fig. S6a, b). To address this problem, we developed a water-in-oil sphere technique to create thin layers of matrix materials over the organoids (Fig. 4b). While embedding organoids in gel domes and layers has been well documented, the large gel volumes of such approaches greatly reduce metabolite transfer owing to the reduced diffusion through a porous hydrogel^16^. Oil-casted hydrogel spheres have multiple advantages: their small volumes allow for efficient exchange of metabolites over traditional approaches; they protect and maintain the tissue architecture of the organoid under shear stresses associated with shaker culture; they provide a basal substrate made of basement membrane components (e.g. MG) or ECM (e.g. collagen) enabling physiological interactions with an ECM; and finally, by mixing other cell types into the precursor gel solutions, they provide a method through which additional cell types can be placed in close proximity with the organoid. With the water-in-oil sphere technique, we were able to produce MG and collagen spheres from as little as 1 µL to 10 µL in volume (Fig. 4c). While organoids could be embedded within MG spheres, supporting large bud formation in the EGM phase (Supplementary Fig. S6c), it was desirable to remove the MG prior to differentiation. This allowed the apicobasal wall to thicken while avoiding significant outgrowth into the surrounding gel; such outgrowth is typical of embedding such organoids within a basement membrane-rich matrix like MG. Therefore, we elected to use dissolved MG in cell medium for the EGM phase and embed organoids in collagen spheres for the MAT phase. We also tested combined MG and collagen spheres for the EGM phase (Days 4-8) but found it inhibited their neuroepithelial expansion (Supplementary Fig. S6d).

**Fig 4.**
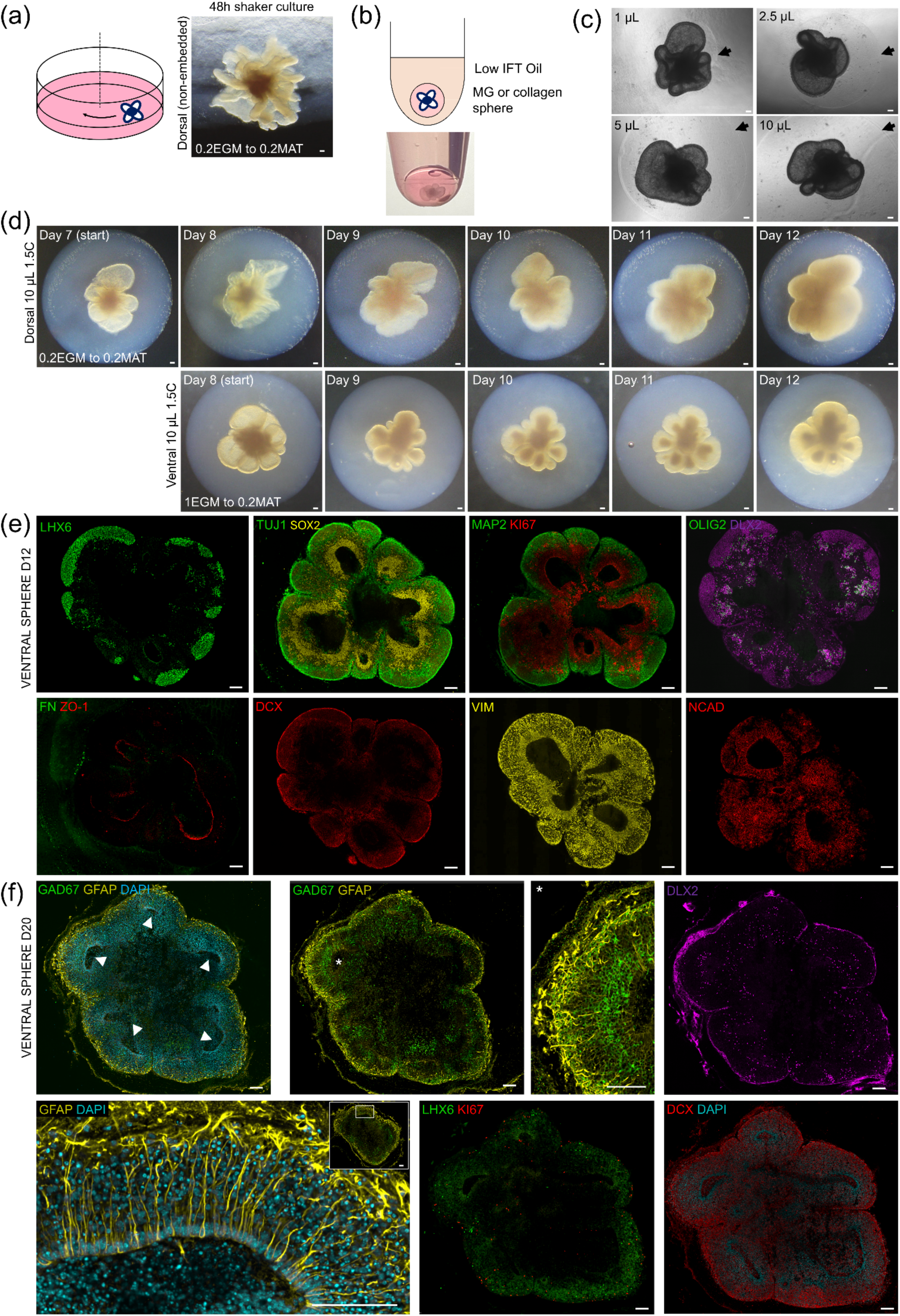
Water-in-oil sphere technique enables small-volume matrix coatings of organoids and preserves tissue architecture under shaker culture. **(a)** Schematic (left) demonstrating shaker culture. Loss of tissue architecture in EGM-cultured organoids when shaker culture is undertaken (right). **(b)** Schematic (top) and image (bottom) of MG-in-oil sphere containing forebrain organoid. Low IFT: low interfacial tension. **(c)** EGM-cultured organoids grown Day 4-8 within miniature MG spheres of 1, 2.5, 5 and 10 µL volumes. Arrows highlight edge of gel spheres. **(d)** Mouse dorsal (top) and ventral (bottom) forebrain organoids embedded in collagen spheres at Day 7 and Day 8 respectively are maintained under shaker culture; preservation of tissue architecture and radial thickening of the neuroepithelium. 1.5C: 1.5 mg/mL collagen hydrogel. **(e)** Collagen-embedded organoids developed under shaker culture show greatly enlarged LHX6+ and DLX2+ regions in contrast to non-embedded, static culture equivalent. TUJ1-SOX2 staining demonstrates clear progenitor and differentiated zones. MAP2-KI67 staining distinguishing postmitotic neurons from proliferative progenitors, with DCX revealing immature neurons. VIM and NCAD markers display structured radial glia scaffolding in ventricular and sub-ventricular zones. **(f)** Extended shaker culture of sphere-embedded mouse ventral forebrain organoids at Day 20. A rich layer of GAD67+ cells is visible in every bud. GFAP displays organised radial glial scaffolding across the apicobasal axis with a rich layer of astrocytes surrounding the organoid. Scalebars: 100 µm.

Using collagen spheres, we were able to maintain tissue architecture for both unguided and ventral (Fig. 4d) ventricular forebrain organoids during shaker culture. This resulted in a thickening of the neuroepithelium following the spatial structural patterning imprinted in the neuroepithelium by the prior EGM-culture. Serial sectioning and immunostaining of the ventral forebrain organoids revealed improved maturation over static culture conditions and non-embedded organoids (Fig. 4e). Notably, more extensive LHX6+ and DLX2+ regions of post-mitotic interneurons and intermediate progenitors, respectively, were detected. Later timepoint staining for GFAP and GAD67 revealed further structure and layering within the bud structure (Fig. 4f). It is worth highlighting therefore that the combination of an expanded neuroepithelium from the EGM medium and the stability provided by the collagen spheres enables the formation of larger and highly organised tissue structures.

### 2.5. Sphere technique enables controlled incorporation of ancillary cell types

The water-in-oil sphere technique also provides a controlled method for delivering a high and uniform density of ancillary cell types to the organoid surface, without compromising organoid viability (Fig. 5a). Using warm coconut oil enabled rapid gelation of collagen hydrogel or MG. This suggested that vascular cells could be incorporated in such a way as to mimic their basal entry point into the developing brain. We therefore co-embedded human umbilical vein endothelial cells (HUVEC) or mouse brain microvascular endothelial cells (MBMEC) alongside mesenchymal stromal cells (MSCs) with organoids in collagen spheres. This localised co-culture led to vascular cell-mediated contraction of the collagen gel spheres (Fig. 5b) and resulted in a cell-rich layer around the organoid, much like the pial membrane in vivo. However, contraction of the 1.5 mg/mL collagen gel sphere also produced a significant densification and alignment of the collagen fibrillar gel, forming a greater barrier to metabolite exchange and a stiffer matrix, which imparted further mechanical stresses upon the organoids and led to a loss of ventricular structure (Fig. 5b).

**Fig 5.**
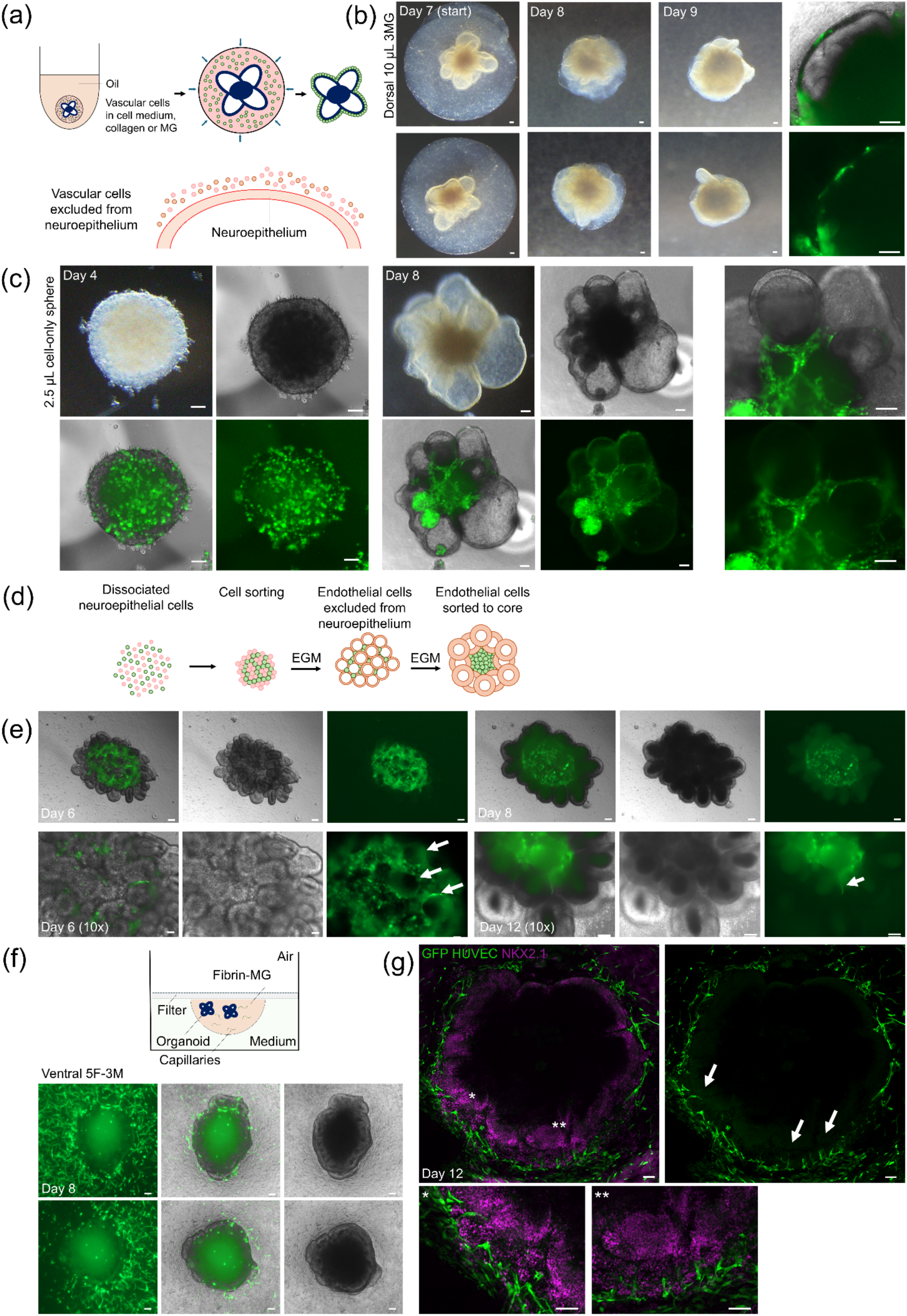
Sphere technique enables controlled incorporation of extra-CNS cell types. **(a)** Schematics showing vascular cell coating technique with ECM sphere or cell suspension sphere approach. **(b)** Vascular cells and organoid co-cultured in collagen spheres for 48h. Cell-mediated collagen gel shrinkage brings GFP-expressing vascular cells into proximity of organoid. **(c)** Cell-only suspension sphere technique applied at Day 4 yields GFP vascular cell coating on forebrain organoids. By Day 8, vascular cells have migrated around basal surface of neuroepithelial buds. **(d)** Schematic depicting endothelial cell incorporation via dissociation technique. **(e)** Time series of dissociated and reaggregated organoids exhibiting endothelial cell exclusion from nascent neuroepithelial structures and cell sorting to core of organoid. Arrows highlight exclusion of GFP vascular cells from neuroepithelial buds. **(f)** Membrane-mounted inverted dome technique; schematic and organoids embedded in fibrin-MG domes alongside GFP vascular cells, Day 4-8. Organoids exhibit limited neuroepithelial expansion when constrained in non-MG substrates. **(g)** When neuroepithelial surface was disrupted by the resultant gel contraction, GFP vascular cells could invade the organoid; * and ** provide magnified examples of this. Scalebars: 100 µm.

We therefore used the sphere technique to deliver just vascular cells to the organoid surface in the absence of matrix materials (Fig. 5c). Despite the proximity and patterning of vascular cells on the organoids, there was a clear exclusion of green fluorescent protein (GFP) vascular cells from the neuroepithelium itself. To further examine the propensity for vascular cells to be excluded from neuroepithelial buds, we dissociated organoids at the post-neural induction stage and mixed in endothelial cells, re-forming a 3D cell aggregate (Fig. 5d). EGM culture from Day 4-8 of the re-aggregated neuroepithelial cells produced numerous small buds, notably still excluding endothelial cells to the periphery of the epithelium (Fig. 5e). Over time, endothelial cells appeared to be sorted to the core of the organoid where a lack of oxygen and nutrients may have led to necrotic cell loss.

### 2.6. Membrane-mounted inverted domes for forebrain organoid vascularisation

We sought to stimulate vascular ingrowth by embedding ventral forebrain organoids and vascular cells in gel domes with VEGF supplementation to the medium (Fig. 5f). By adding organoids post-neural induction (Day 4), alongside vascular cells, to fibrin-MG domes, we observed characteristic capillary formation adjacent to the organoid. Fibrin was used as a pro-angiogenic matrix which has a rapid gelation time compared to collagen or MG, thus enabling the vascular cells to be embedded uniformly in the gel. However, the presence of fibrin limited ventricular expansion typical of EGM-culture. In addition, despite the presence of VEGF in the media, no capillaries were observed to invade into the organoid. Embedding vascular cells in collagen gel domes at Day 8, we observed sprouting into the periphery of the organoid (Fig. 5g). However, this was limited by cell-mediated gel contraction as the gel had shrunk up onto the supporting membrane and begun to disrupt the neuroepithelium, allowing endothelial cell invasion. Therefore, where we have observed examples of vascularisation, it appeared to be due to a disruption of the architecture of the neuroepithelial boundary.

### 2.7. Expanded ventricular forebrain organoids reveal challenges to developing vascularised embryonic tissue architecture

In the mouse ventral forebrain model, we observed a contiguous, outer layer of DCX+ and LHX6+ cells, parallel to the ventricular surface (Fig. 6a), as reported in the mouse brain^17^, formed through the apicobasal thickening of the neuroepithelial layer. The LHX6+ region stopped where individual buds contacted, suggesting that the EGM-enlarged ventricles provide sufficient space, and transport of oxygen and nutrients, for higher order cell layered architecture to form which is comparable to the *in vivo* MGE. In typical organoid culture, neurons mature around many, miniature ventricles contained within the aggregate body, or form a band of mature cells around the edge of the organoid, which is representative of the clear apicobasal thickening of the neuroepithelial layer which occurs *in vivo*. As expected, the EGM-derived organoids displayed markers for radial glial scaffolding (vimentin and N-cadherin) once differentiation had taken place by Day 12 (Fig. 6b). Previous research suggests that radial glia can provide either a direct scaffold for vascular ingress or provide paracrine signals (e.g. VEGF) to this end, thereby playing a key role in vascularisation dependant on tissue architecture^18^. We found the radial glia in the ventral forebrain organoids were more disorganised in the upper layers than around the ventricle as determined by vimentin and N-cadherin immunostaining.

**Fig 6.**
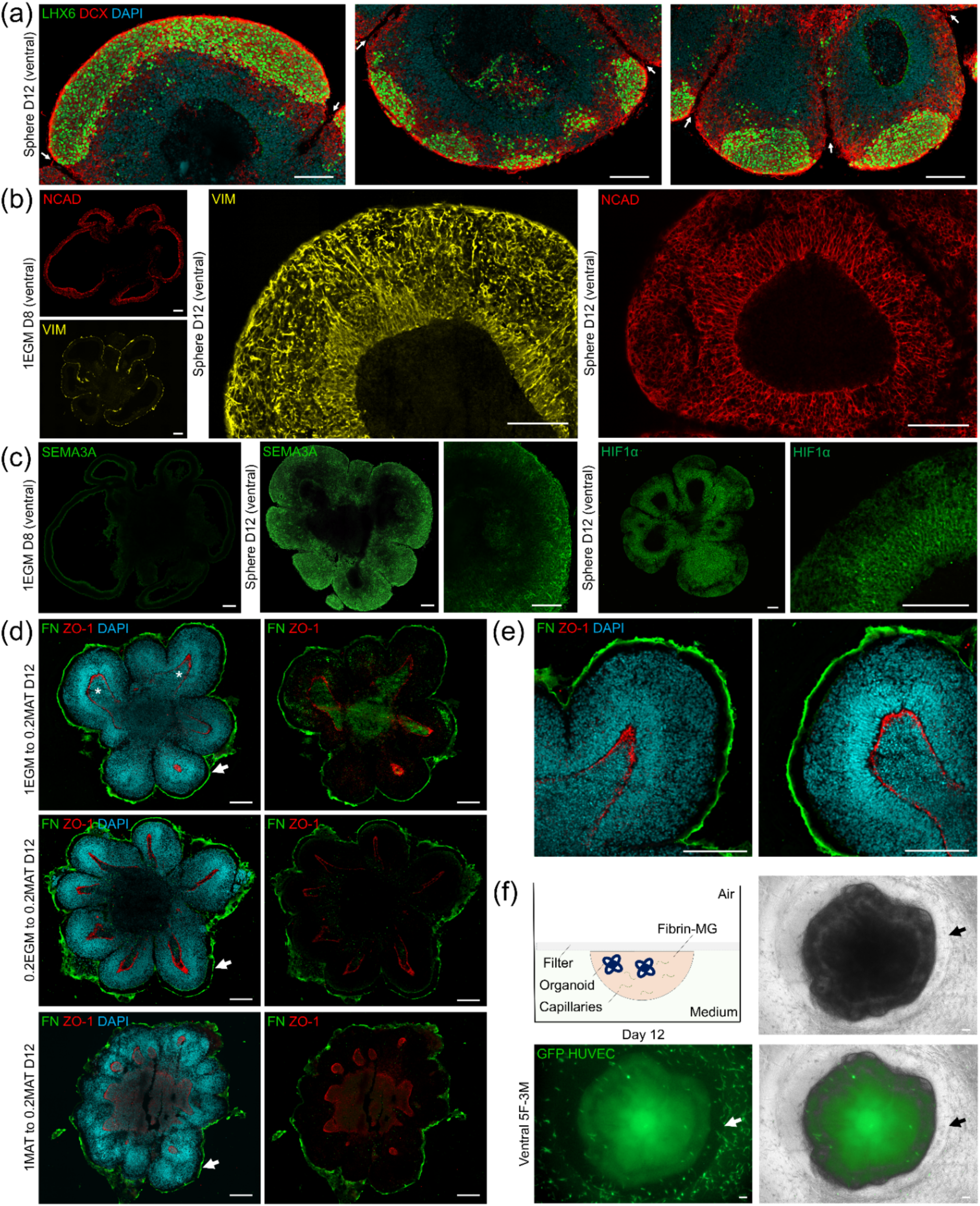
Large ventricular forebrain organoids reveal challenges to developing embryonic tissue architecture. **(a)** EGM-enlarged ventricles provide sufficient space for a large, laminar MGE-like regions of LHX6+ and DCX+ cells in mouse ventral forebrain organoids. No LHX6+ signal was detected in DCX+ regions between buds. Arrows highlight individual bud boundaries. **(b)** Disorganised radial glia scaffolding outside of ventricular and sub-ventricular zones as reported by vimentin and N-cadherin markers. **(c)** SEMA3A present in Day 12 organoid may restrict vascularisation in neurogenic organoids. HIF-1α detected in the peri-ventricular layers – localised in the cytoplasm, but not nucleus – comprising neural stem cells and progenitors. **(d)** Loss of basement membrane adhesion to forebrain organoids for different Day 4-8 conditions: 1EGM; 0.2EGM; and 1MAT. **(e)** Magnified 1EGM to 0.2MAT buds highlighting detachment of FN layer from organoid bud. No nuclei were detected in the gap. **(f)** Organoids embedded within fibrin-MG matrix exhibited protease-driven degradation of matrix around organoid (highlighted by arrows); similar degradation was not observed within collagen spheres. Scalebars: 100 µm.

Semaphorins, notably SEMA3A, have been identified as important components not only for axonal guidance but also brain vascular patterning, regulating and restricting angiogenesis from the surrounding pia^19–21^. We detected negligible SEMA3A in the post-EGM day 8 organoids (Fig. 6c), whereas following Days 8-12 differentiation in MAT medium, we detected Sema3A in the outer neuronal regions of the organoid. Simultaneously, we observed negligible Hif-1α signal in Day 8 pre-differentiation (Supplementary Fig. S2a) and HIF-1α+ regions in Day 12 samples in the ventricular zone, in the cytoplasm of progenitor cells, but distinctly lacking from the outer neuronal layers (Fig. 6c). The continuous expression of HIF-1α by neural stem cells and progenitors and its localisation in the cytoplasm, but not nucleus, is expected under normoxic conditions and plays a role in physiological neural progenitor function^22^. This suggests that important aspects of in vivo development of receptive neural compartments can be recapitulated with the EGM protocol. However, the narrow timing window between radial glia formation – with associated pro-angiogenic signalling – and neurogenic layer formation – with anti-angiogenic SEMA3A expression – may explain why we do not see angiogenesis into the neurogenic layer with the current protocol.

While we observed that fibronectin was highly expressed on the basal surface of the EGM-cultured organoids by Day 8, we observed a clear gap between the basal fibronectin and the basal surface of the organoid buds at Day 12 (Fig. 6d, e). Organoids grown in EGM or MAT medium with 1 mg/mL GFR MG for days 4-8 were removed of Matrigel, which may have explained the detachment; however, organoids grown in EGM with 0.2 mg/mL GFR MG also exhibited this property. We also embedded Day 8 organoids in an interpenetrating gel of 5 mg/mL fibrin and 3 mg/mL MG and observed a degradation of the fibrin gel around the organoid (Fig. 6f, Supplementary Fig. S6e). The detection of fibrinolysis here, while not detecting similar degradation within collagen sphere-embedded organoids, is suggestive of serine proteases (likely plasmin). Plasmin is less effective at degrading collagen but, interestingly, is effective at degrading fibronectin, which at low or mild levels affects cell-fibronectin integrin binding, and may therefore explain the basal layer detachment of Fig. 6e^23–26^, and which may also hinder vascular ingress across this surface. Although challenges remain, the combination of EGM-induced expansion and cell-impregnated hydrogels yields better modelling of neurogenic neuroepithelium, its response to ECM, and interaction with non-neural cells. The current methodology is a promising basis for further investigation that may leverage the improved neuroepithelial architecture to achieve vascularisation.

### 2.8. Human EGM-cultured organoids produce greatly enlarged unobstructed ventricular lumens and delayed maturation

Finally, we applied the EGM culture method to human iPS cells and adjusted stage timings to match the established timing of neural induction for human cells (Fig. 7a), which revealed that by Day 19, EGM produced buds with an well-established neuroepithelial monolayer, enclosing an unobstructed ventricular lumen, for ventral and dorsal human organoids (Fig. 7b). Overall organoid size was not significantly different between EGM and MAT-derived human organoids, at least by Day 19 (Fig. 7c – top), and bud number was only significantly different for the dorsal variation (Fig. 7c – bottom). However, MAT-derived human organoids did not produce unobstructed ventricular lumens distinct from the central organoid mass.

**Fig 7.**
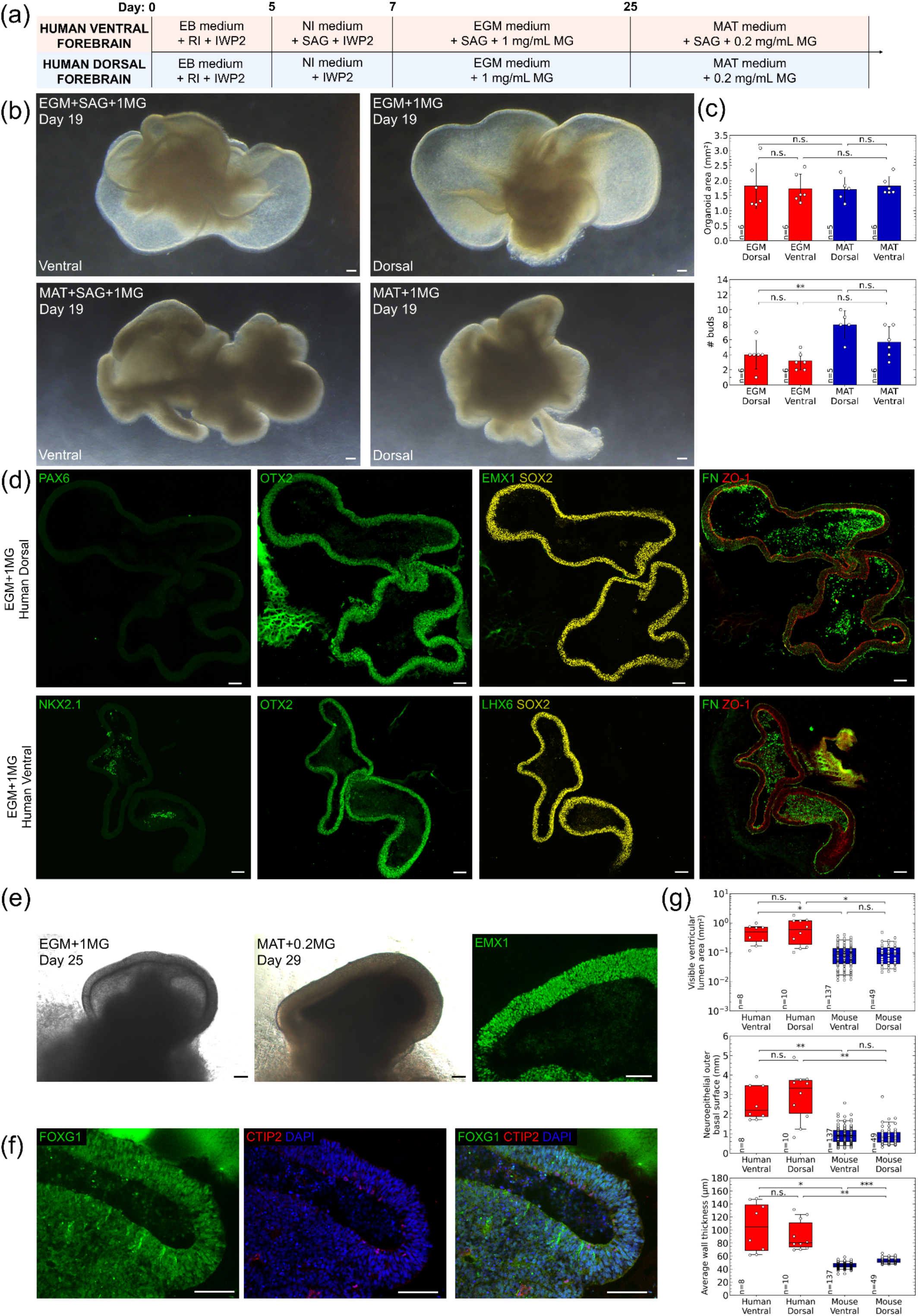
Human brain organoids exhibit neuroepithelial expansion and delayed regional commitment. **(a)** Protocol for human ventral and dorsal forebrain organoids. **(b)** Neuroepithelial expansion with EGM for ventral and dorsal human forebrain organoids, compared to those cultured in MAT medium. **(c)** Measurements of human organoid area and bud number show no significant difference between EGM and MAT culture conditions at Day 19. **(d)** Organoids cultured in EGM and MAT mediums from Day 7-19 are positive for anterior neuroectoderm marker (OTX2). **(e)** Human dorsal organoids cultured in EGM to Day 25 (left) were moved to MAT medium until Day 29. By Day 29 (centre), organoids exhibit apicobasal thickening, with EMX+ marker throughout (right). **(f)** FOXG1 staining demonstrates acquisition of forebrain fate, while negative CTIP2 marker shows no neurons were present. **(g)** Measurements of ventral and dorsal human organoids derived using EGM: visible ventricular lumen area (top); length of outer basal surface in cross-section (middle); average neuroepithelial wall thickness (bottom). Statistics: Levene’s test followed by Kruskal–Wallis with Games–Howell post-hoc comparisons. p < 0.05 (*), p < 0.01 (**) and p < 0.001 (***); n.s. means no statistically significant difference between groups was detected. Scalebars: 100 µm.

Immunostaining of the EGM-cultured human organoids at Day 19 (Fig. 7c) showed no differentiation of the neuroepithelial layer by Day 19 (SOX2+ throughout the neuroepithelial layer, EMX1-/LHX6-) and apicobasal polarisation (FN/ZO-1). Regional identity markers (PAX6, NKX2.1) were negative in organoids cultured continuously in EGM, instead expressing the transcription factor OTX2, which marks the entire anterior neuroectoderm before becoming more restricted to the diencephalon and mesencephalon. MAT-cultured organoids displayed some regions positive for PAX6 (dorsal) and NKX2.1 (ventral) (Supplementary Fig. 7). Human organoids moved from EGM to a maturation medium at Day 25, began to show by Day 29 an apicobasal thickening of the neuroepithelium (Fig. 7d – left, centre), and positive staining for forebrain markers EMX1 (Fig. 7d – right) and FOXG1 (Fig. 7e), but still did not show production of neurons even at this late time point (CTIP2-), demonstrating a much later acquisition of regional fate compared with mouse organoids. By comparing the geometric features of human and mouse organoids, we observed species-specific differences in unobstructed ventricular lumen area (Fig. 7f – top), neuroepithelial outer basal surface (Fig. 7f – middle), and average wall thickness (Fig. 7f – bottom). While mouse organoid ventricular lumens did not expand after 4 days of continual EGM treatment, instead leading to thickening of the neuroepithelial wall, human organoid ventricular lumens continued to grow up to at least Day 29, leading to significantly larger organoids while the neuroepithelium remained relatively thin. Thus, EGM culture appears to enable cell intrinsic, species-specific capacity for expansion and delay in maturation – otherwise accelerated by standard brain organoid culture – producing expanded human ventricular lumens reminiscent of the in vivo developing first trimester brain.

## 3. DISCUSSION

Organ geometry influences progenitor pool size, microscale tissue architecture and layering, organiser diffusive gradients, and the interaction with extrinsic neurovascular cell types. Without the early-stage pre-requisites for tissue structure, organoid models have been limited in attempts to generate higher-order organisation, therefore restricting their physiological temporal development. Further, a lack of proper tissue geometry and structure may be linked to the key challenge of vascularising brain organoids. While several studies have demonstrated endothelial and supportive cell types within organoids, ingression into the neuroepithelium as *in vivo* has, to date, not been concretely demonstrated *in vitro*.

EGM culture medium is supplemented with several mitogenic growth factors – bFGF, EGF, IGF – which are known to drive neuroepithelial cell proliferation, while other components – ascorbic acid, hydrocortisone, and serum – reduce oxidative stress. However, these supplements alone, when added to standard neurogenic mediums, did not produce large ventricular budding. Instead, we found that the basal composition of endothelial cell mediums was central to the expansion, maintaining a stem-like state (SOX2+) and promoting tangential expansion of the ventricle. The subsequent culture in a neural maturation medium enabled apicobasal thickening and differentiation to a layered forebrain tissue with proper cytoarchitecture.

We propose that the basal medium stabilizes the glycolytic metabolic program already present in neural stem cells and progenitors, even under normoxic conditions, by maintaining a specific intracellular reactive oxygen species (ROS) balance, through metal and trace elements. ROS signalling (e.g. via NRF2 and NFkB) has significant effects on stem cell behaviour, with relatively low ROS promoting self-renewal, proliferation, and suppression of differentiation signals^27–29^. Consistent with this metabolic state, neural stem cells preferentially rely on glycolysis while maintaining comparatively low mitochondrial oxidative metabolism^30^. In contrast, elevated ROS levels have been reported to promote exit from the stem cell state and trigger neuronal differentiation^31^. EGM culture may find application in other organoid tissue-types, in which it may be used as a useful intermediate stage between lineage determination and differentiation, enabling tissue-scale architecture within organoid culture systems.

Vegetable oils provide a biocompatible, hydrophobic fluid for forming miniature, aqueous-based gel spheres via a water-in-oil drop technique as demonstrated in this study. Of refined vegetable oils, coconut oil has the lowest interfacial surface tension (13.9 mN/m) while others, such as sunflower oil, have a notably higher interfacial surface tension (24.8 mN/m)^32^. The importance of this lies in the removal of an oil film from the sphere when re-immersed in cell medium; an oil barrier otherwise limits the exchange of metabolites between the aqueous-filled pores of the gel and aqueous cell medium. Tissue biomechanics has a significant influence upon neural development with many mechanical cues arising from tissue architecture. This includes matrix stiffness from the surrounding ECM, which also plays a key role in angiogenesis; cytoskeletal forces; and hydrostatic pressure and flow-derived shear stresses from the ventricular fluid (i.e. cerebrospinal fluid)^33,34^. Furthermore, the compliance of the ECM substrate has been reported to influence the selection of neuronal vs glial differentiation pathways^35^. Therefore, alongside the maintenance of the ventricular lumen via the EGM culture approach, we developed a method to embed organoids in small volume collagen gel spheres. The Elastic modulus of the brain is reported to be 0.1-1 kPa^36^; 1 and 2 mg/mL collagen gels are reported to have elastic moduli of 170 Pa and 328 Pa, respectively^37^, and 1.5 mg/mL collagen gels to have a Young’s Modulus of 100 Pa^38^ from which we chose a 1.5 mg/mL collagen gel matrix likely with an Elastic modulus in the range of 250 Pa. When coupled with the EGM-expanded organoids, the result was a layered telencephalic structure and no migration of axons into the gel. Further investigations could study how the tuneable hydrogel sphere stiffness and composition can affect brain organoid architecture.

There has been significant interest recently in vascularising brain organoids given the central importance that vascularisation plays in the function, maintenance and pathology of neural tissues. Several groups report the invasion or fusing of vascular cells (or vascular organoids) into cerebral organoids^39^. In contrast, we observed no invasive vascularisation, except in cases where the neuroepithelial layer was disrupted. In standard brain organoid protocols, multiple neuroepithelial buds are present within a single organoid and the basal surface apposition between buds may provide access for vascular cells to enter between buds without invading the thickening neuroepithelium. Ingression into the neuroepithelium, as in vivo, is thus still lacking, and we hope this work provides a suitable model for future work studying vascular invasion from which one can easily discriminate vascular invasion into the neuroepithelium versus migration between neuroepithelial buds. Vascularisation first occurs through the ventral forebrain which, rather than a passive response to oxygen and nutrient demands, follows a highly programmatic patterning process via region-specific homeobox transcription factors^40^ and thus likely requires considerable tissue-scale patterning to achieve this objective.

In this paper, we have reported a new approach to generating and maintaining tissue-scale architecture within forebrain organoids. Through EGM-mediated culture, forebrain organoids exhibit expansion of the undifferentiated neuroepithelium, forming a thin-walled structure reminiscent of the embryonic brain vesicle. The support of an even more extended timeline and greater enlargement in human organoids mirrors the prolonged human developmental timeline compared with mouse and supports the notion that this delay is responsible for the extended size. Interestingly, mouse organoids even when continually cultured in EGM for longer could not further expand, suggesting a species-specific difference in intrinsic metabolic requirements and capacity to expand. Thus, this approach provides a tool to investigate neurodevelopmental principles, while the incorporation of supporting vascular cells will enable further studies concerning vascular ingress into the developing brain.

## METHODS

### Reagent preparation

Embryoid body (EB), neural induction (NI), and Maturation (MAT) medium were prepared from STEMdiff Cerebral Organoid Kit (STEMCELL Technologies) and supplemented as according to provider protocols. Endothelial cell growth medium 2 (EGM, Promocell) was supplemented using the kit without heparin. Vascular endothelial growth factor 165 (VEGF165) and basic fibroblast growth factor (bFGF146, Peprotech) were resuspended at 100 µg/mL in phosphate-buffered saline (PBS) containing 0.1 %(w/v) human serum albumin (HSA) and supplemented into cell medium at 40 ng/mL. Supplement components of EGM were added excluding heparin at concentrations based on EGM composition: 2%(v/v) fetal calf serum, 5 ng/mL epidermal growth factor (EGF, Peprotech), 0.5 ng/mL VEGF165, 10 ng/mL bFGF, 20 ng/mL insulin-like growth factor (IGF-R3L), 0.2 µg/mL hydrocortisone (STEMCELL Technologies) and 1 µg/mL ascorbic acid. Medium 200 and MCDB 131 (both Thermo Fisher Scientific) were similarly supplemented as described. Smoothened agonist (SAG, Merck) was resuspended at 1 mM in water and supplemented into cell medium at 100 nM. IWP2 was resuspended in DMSO at 5 mM stock concentration and supplemented into cell medium at 2.5 µM. CHIR-99021 was resuspended in DMSO at 10 mM, PD03259010 was resuspended in DMSO at 1 mM, mLif was resuspended in PBS containing 0.1 %(w/v) bovine serum albumin (BSA) at 10 µg/mL and Y-27632 Rock inhibitor (RI) was resuspended in water at 5 mM. Matrigel (MG) and Growth factor reduced Matrigel (GFR MG) (Corning) were thawed on ice and stock concentrations were recorded based on Lot number. Fibrinogen (Merck) was resuspended at 25 mg/mL in 0.9 %(w/v) NaCl in water at 37 °C for 2h. Thrombin (Merck) was resuspended at 50 U/mL in 0.9 %(w/v) NaCl + 0.1 %(w/v) HSA in water. SFES+2iLif medium: 50% DMEM/F12 (Gibco), 50% Neurobasal medium (Gibco), 1:200 N2 supplement (Gibco), 1:100 B27 supplement (Life Technologies), 1:150 BSA solution and 1:100 penicillin-streptomycin (PS), supplemented with 3 µM CHIR-99021, 1 µM PD03259010, 10 ng/mL mLif, and 1:500 RI.

### Cell lines

GFR MG-coated 6 well plates were prepared at 8.7 µg/cm^2^ and gelled for 1h prior to use. Naive mouse E14 ESCs were thawed and initially plated onto GFR MG-coated 6 well plates using SFES+2iLif cell medium. After 3 days, cell medium was changed for StemFlex medium supplemented with 10 ng/mL mLif, for priming. Following priming, E14s were routinely subcultured using EDTA for 5 min or passaged using TrypLE Express for 5 min, followed by centrifugation (250 g, 5 min) and counting. Human male iPSC line BIONi010-B (Sigma, EBiSC) was cultured on GFR MG-coated 6 well plates using StemFlex cell medium. GFP mouse brain microvascular endothelial cells (C57BL/6-GFP, MBMEC, 2B Scientific), GFP human umbilical vein endothelial cells (GFP HUVEC, CCD-6011, Caltag Medsystems) and bone marrow mesenchymal stromal cells (MSC, Promocell) used at passage 4 or 5 using cell-specific proprietary cell mediums (Promocell).

### Organoid generation

E14s were passaged, centrifuged and counted and transferred to a 96 ultra-low attachment (ULA) well plate (Corning) and cultured in EB medium supplemented with IWP2 for 2 days, and were then incubated in NI medium for a further 2 days. 32,000 cells were seeded per well for mouse organoids and 16,000 cells for human owing to differences in intrinsic cell timing. For ventral organoids, NI was supplemented with 100 nM smoothened agonist (SAG). Organoids were subsequently transferred in 24 ULA well plates containing EGM2 (Promocell) supplemented with 0.2 or 1 mg/mL GFR MG for 3-4 days. Organoids cultured using 1 mg/mL GFR MG were incubated in Cell Recovery Solution (Corning) for 15-20 min on ice to remove MG. Organoids were then either embedded within collagen spheres (see below) or left non-embedded, and cultured in MAT medium containing 0.2 mg/mL GFR MG.

### Water in-oil collagen hydrogel sphere generation

Collagen gel precursor solutions were prepared following an established method^41^. Briefly, the precursor solution contained 10 % of 10X PBS, 3 % of 7.5 % sodium bicarbonate, 1.5 mg/mL collagen from a 10 mg/mL collagen stock solution (Cell Systems), and EGM or MAT medium. For embedding cells and organoids within collagen spheres, collagen was neutralised in precursor solution, prior to adding cells and organoids. Coconut oil was melted at 37 **°**C. The cell-laden collagen precursor solution was then carefully pipetted using a cut P10 tip, pre-washed with 5% bovine serum albumin (BSA) to prevent organoid attachment, to a 1.5 mL Eppendorf tube containing warm coconut oil and incubated at 37 **°**C for 30 min. The pre-warmed coconut oil leads to rapid gelation of the collagen precursor solution. The gelled sphere was subsequently removed by replacing most of the oil with warm cell medium, injected close to the sphere to break the surface tension with the oil. Once floating in the cell medium, the sphere was transferred to a well plate using a cut P1000 tip, pre-washed with 5% BSA.

### Dissociated organoid protocol

Organoids were dissociated at Day 4 using TrypLE for 5 min and subsequent repeated pipetting. Organoid-derived cells were subsequently mixed with vascular cells and centrifuged at 250 g for 5 min within 96ULA well plates, and EGM supplemented with 1 mg/mL GFR MG was added. After 24 h, cell aggregates were moved to 24ULA well plates.

### Inverted dome protocol

Fibrin-MG domes were created by preparing precursor solutions comprising 5 mg/mL fibrinogen and 3 mg/mL GFR MG. Thrombin was added at 1 U/mL to induce gelation to fibrin. 25 µL gel domes were formed on well plate inserts which were placed upside-down into 6 well plates. Following 30 min incubation at 37 **°**C, medium was pipetted around the edges of the membrane to immerse the gel dome.

### Organoid processing for immunohistochemistry

Samples were fixed in 10 % formalin for 24 h and subsequently washed three times using PBS. Samples were incubated for 24 h in phosphate buffer (PB) containing 30 % sucrose, as a cryoprotectant, containing a few drops of FastGreen FCF (Sigma). Samples were then embedded in 7.5 % gelatin containing 10 % sucrose in PB and rapidly frozen at -50 **°**C using an isopentane bath for 1 min. Samples were subsequently cryosectioned using a Leica CM1950 and stored at -20 **°**C prior to staining.

### Immunohistochemistry

Slides were immersed in PBS in 50 mL Falcon tubes for 1-2 h at 37 **°**C to remove mounting gelatin. For permeabilization, sections incubated in 0.25 %(v/v) Triton X-100 in PBS for 10 min and subsequently blocked using 0.1 %(v/v) Triton X-100 in 1 %(w/v) BSA and 4 % normal donkey serum (NDS). Primary antibodies were incubated overnight at 4 **°**C in blocking buffer using dilutions found in Supplementary Fig. S8. Sections were washed three times using 1 %(w/v) BSA in PBS and incubated with secondary antibodies for 2 h at room temperature, which were added at 1:500 in blocking buffer. Following three further washes with 1 %(w/v) BSA in PBS, sections were quenched for autofluorescence using TrueVIEW (VectorLabs) for 5 min and washed using PBS. Sections were counterstained with DAPI at 1:500 in PBS for 10 min and mounted using Vibrance Antifade.

### Imaging

Transmission microscopy was undertaken using an EVOS microscope (Thermo Fisher Scientific). Immunofluorescent imaging was conducted using an Andor BC43 (Oxford Instruments). Images were prepared using Imaris.

### Statistical Analyses

In all figures, data are expressed as the arithmetic mean ± standard deviation over *n* independent replicates. Individual data points are plotted where appropriate. Levene’s tests were used to determine whether the variances were homogenous. Variances were found to not be homogenous, therefore data were analysed for statistical significance using the Kruskal-Wallis and Games-Howell pairwise post hoc tests. Differences were considered statistically significant at *p* < 0.05 (*), p < 0.01 (**) and p < 0.001 (***); n.s. means no statistically significant difference between groups was detected.

## AUTHOR CONTRIBUTIONS

AWJ and MAL conceived of and designed the research. AWJ performed experiments and wrote the manuscript. AA and LG helped with organoid generation and characterisation. MAL supervised the work and acquired funding. All authors provided critical revision and final approval of the manuscript.

## ACKNOWLEDGEMENTS

The authors would like to thank members of the Lancaster lab for feedback and technical support. This work was supported by funding from UKRI Medical Research Council (MC_UP_1201/9) and a Vallee Scholars Award.

## SUPPLEMENTARY FIGURES

**Supplementary Figure S1.**
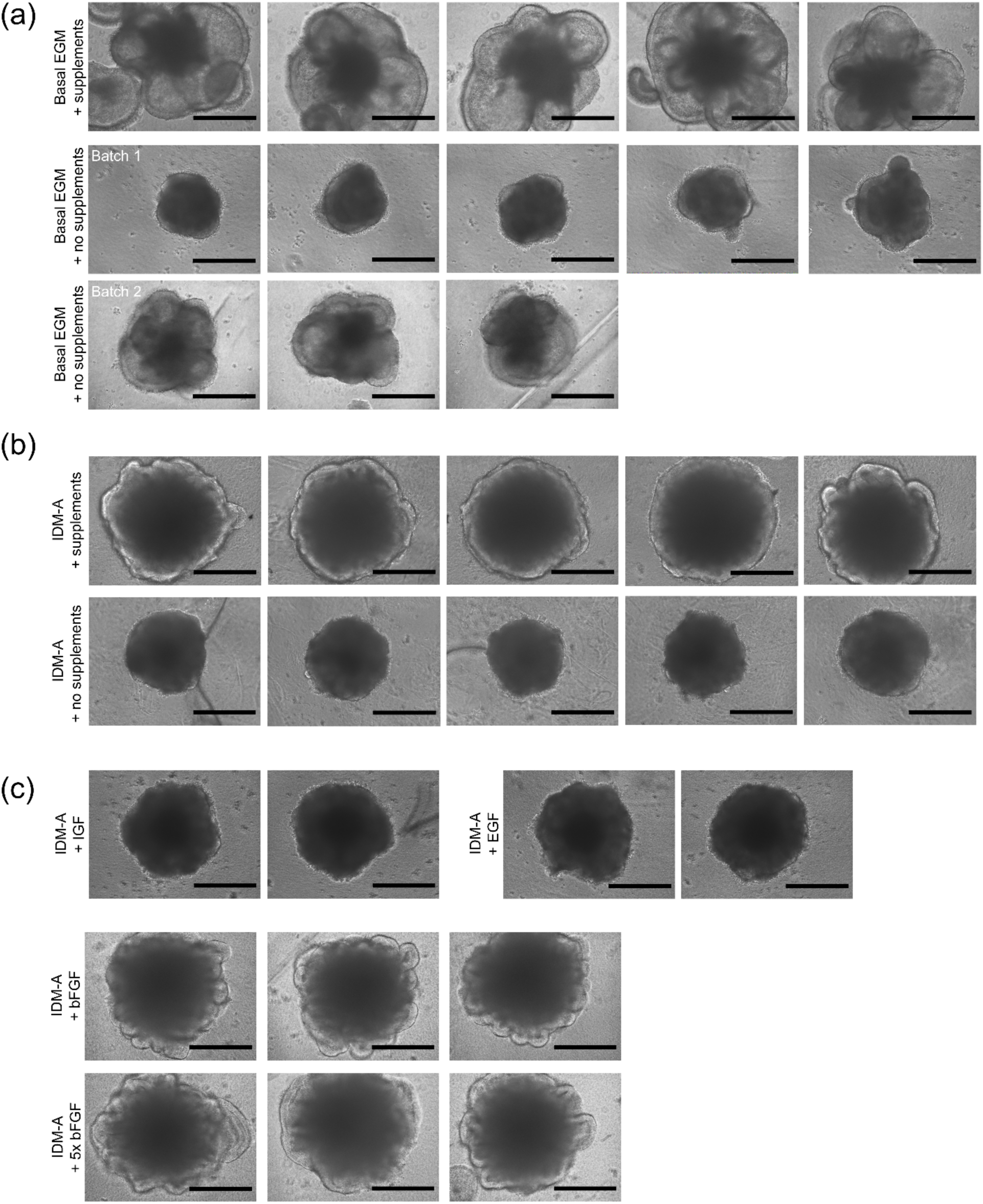
Effect of growth factor supplementation of EGM and IDM mediums on bud growth. **(a)** Comparison of EGM with and without growth factor supplements shows bud formation even without supplementation. **(b)** IDM-A with EGM supplements shows enlarged organoids organoids but without characteristic bud expansion. (c) IDM-A with individual growth factors: IGF, EGF and bFGF. bFGF shows bud growth but not the level of EGM culture. Scalebars: 400 µm.

**Supplementary Table 1.** . HIF and hypoxia-inducing components within endothelial cell mediums.

| Medium component | Effect |
| --- | --- |
| Ammonium Molybdate | Ammonium molybdate forms a complex with H<sub>2</sub>O<sub>2</sub>, protecting it from catalase activity. Catalase activity produces O<sub>2</sub> and destabilises HIF-1α.<sup>1-5</sup> |
| Ammonium Metavanadate | Vanadate induces HIF-1α expression through ROS (H<sub>2</sub>O<sub>2</sub>)<sup>6</sup> |
| Manganese Sulfate | Manganese inhibits HIF-1α-prolyl hydroxylase, the key enzyme for HIF-1α hydroxylation and subsequent von Hippel-Lindau (VHL)-dependent HIF-1α degradation.<sup>7</sup> |
| Nickelous Chloride | Nickel activates HIF-1α by substituting for Fe<sup>2+</sup> in the regulatory dioxygenases and this substitution inactivates the enzymes.<sup>8</sup> |
| Sodium Meta Silicate | Silicate ions stabilize HIF-1α expression by blocking prolyl hydroxylase domain (PHD2) enzyme.<sup>9</sup> |
| Stannous Chloride | Metal-induced hypoxia through ROS generation.<sup>10</sup> |
| Adenine | HIF-1α stabilisation in already hypoxic cells.<sup>11</sup> Extracellular adenosine levels increase due to extracellular catabolism of released adenine nucleotides. Adenosine can increase HIF-1α protein accumulation.<sup>12</sup> |

**Supplementary Figure S2.**
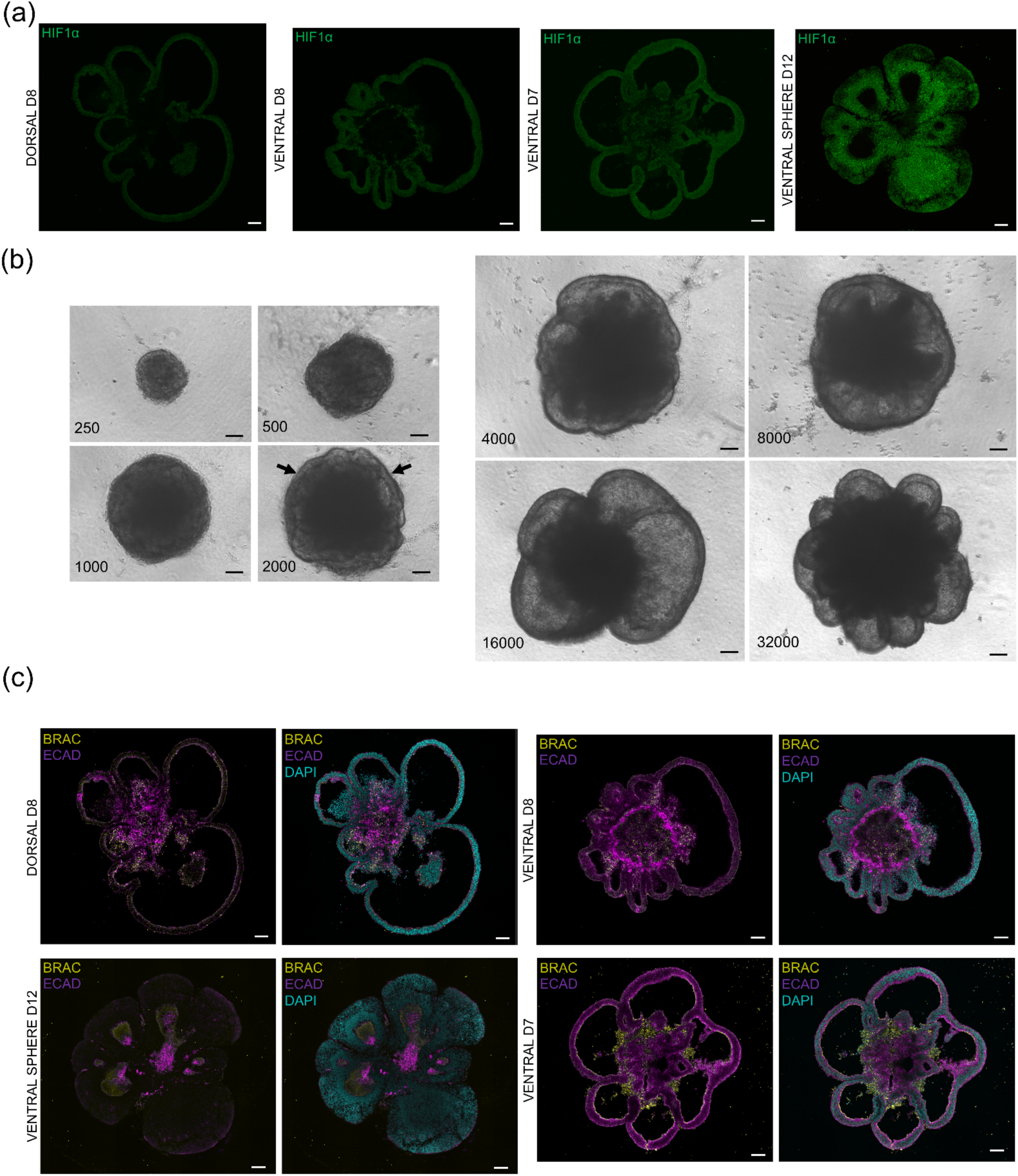
Additional characterisation of EGM-cultured organoids. **(a)** HIF-1a immunostaining for dorsal and ventral organoids at Day 8; ventral organoid at Day 7; and ventral sphere at Day 12, corresponding to mouse ventral organoid embedded within collagen sphere. **(b)** Expanded ventricular organoids are dependent upon number of starting cells. Arrows highlight existence of distinct neuroepithelial wall at threshold size. **(c)** Brachyury and E-cadherin immunostaining revealing structure of core attachment. Scalebars: 100 µm.

**Supplementary Figure S3.**
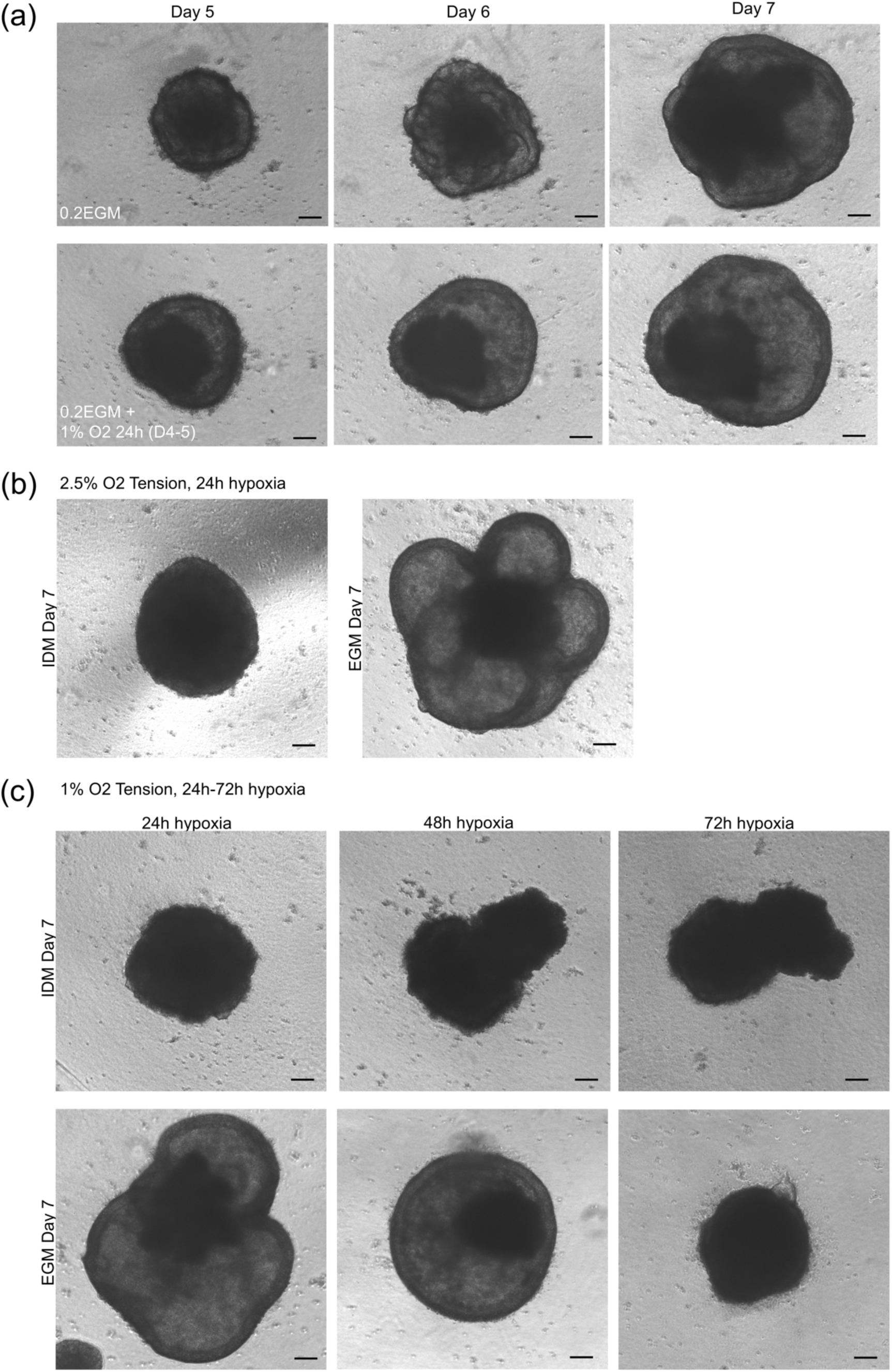
Independence of oxygen tension and EGM upon neuroepithelial expansion. **(a)** Comparison of dorsal organoids (8000 cells/organoid) cultured in EGM + 0.2 mg/ml GFR MG with and without 24h of 1% oxygen tension (Day 4-5). **(b)** Comparison of Improved Differentiation Medium (IDM) to EGM when exposed to 24h of 2.5% oxygen tension. No neuroepithelial expansion was observed in IDM-cultured organoids. (c) Comparison of IDM to EGM-cultured organoids in extended hypoxic conditions. Scalebars: 100 µm.

**Supplementary Figure S4.**
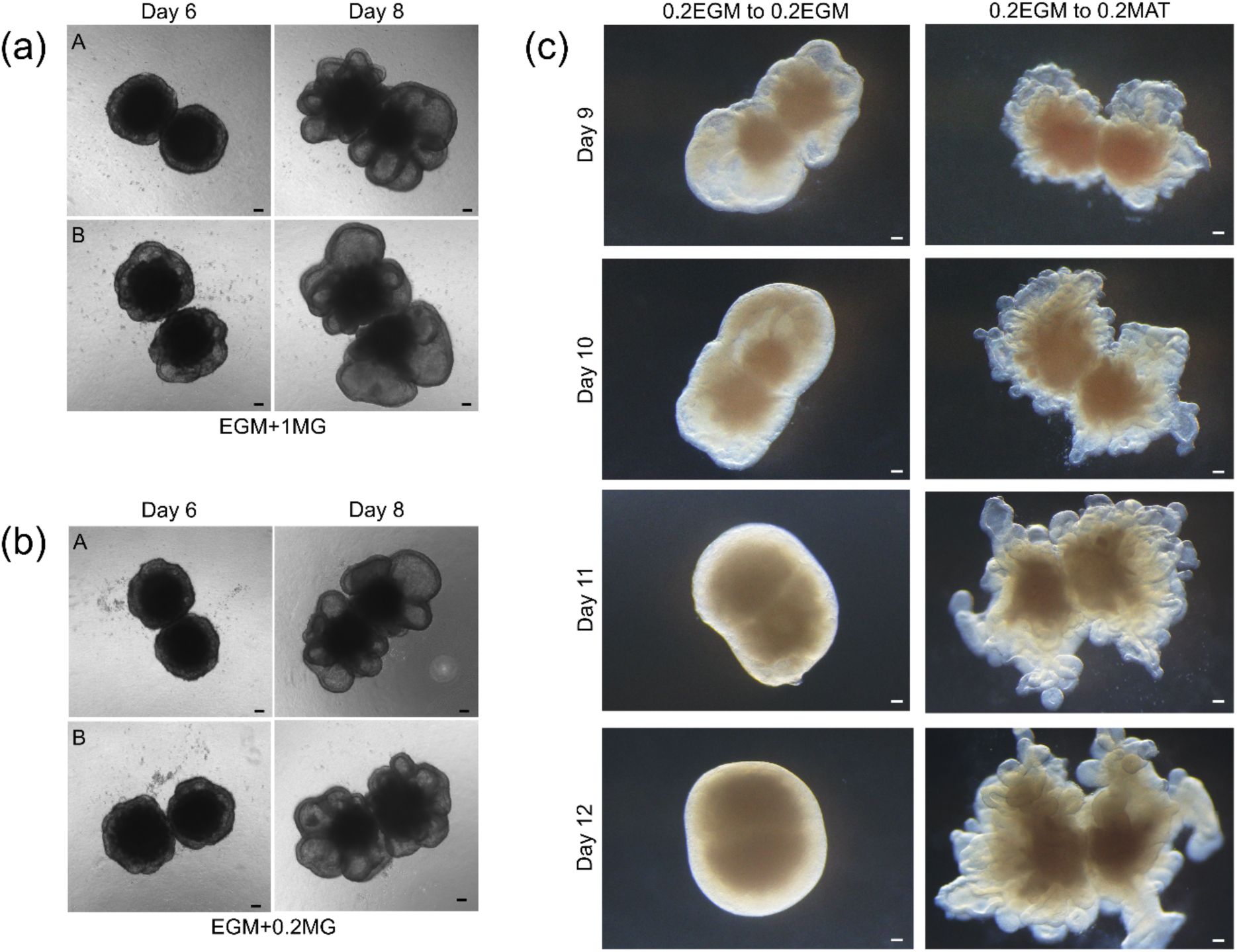
Dorsoventral assembloids using EGM-culture. The expanded neuroepithelial walls of dorsal and ventral organoids cultured together using (a) EGM + 1 mg/ml GFR MG and **(b)** EGM + 0.2 mg/ml GFR MG did not join into single structures, despite organoid mass fusing. (c) 0.2EGM organoids moved to 0.2MAT formed chaotic structures, in contrast to singular ventral and dorsal organoids. Scalebars: 100 µm.

**Supplementary Figure S5.**
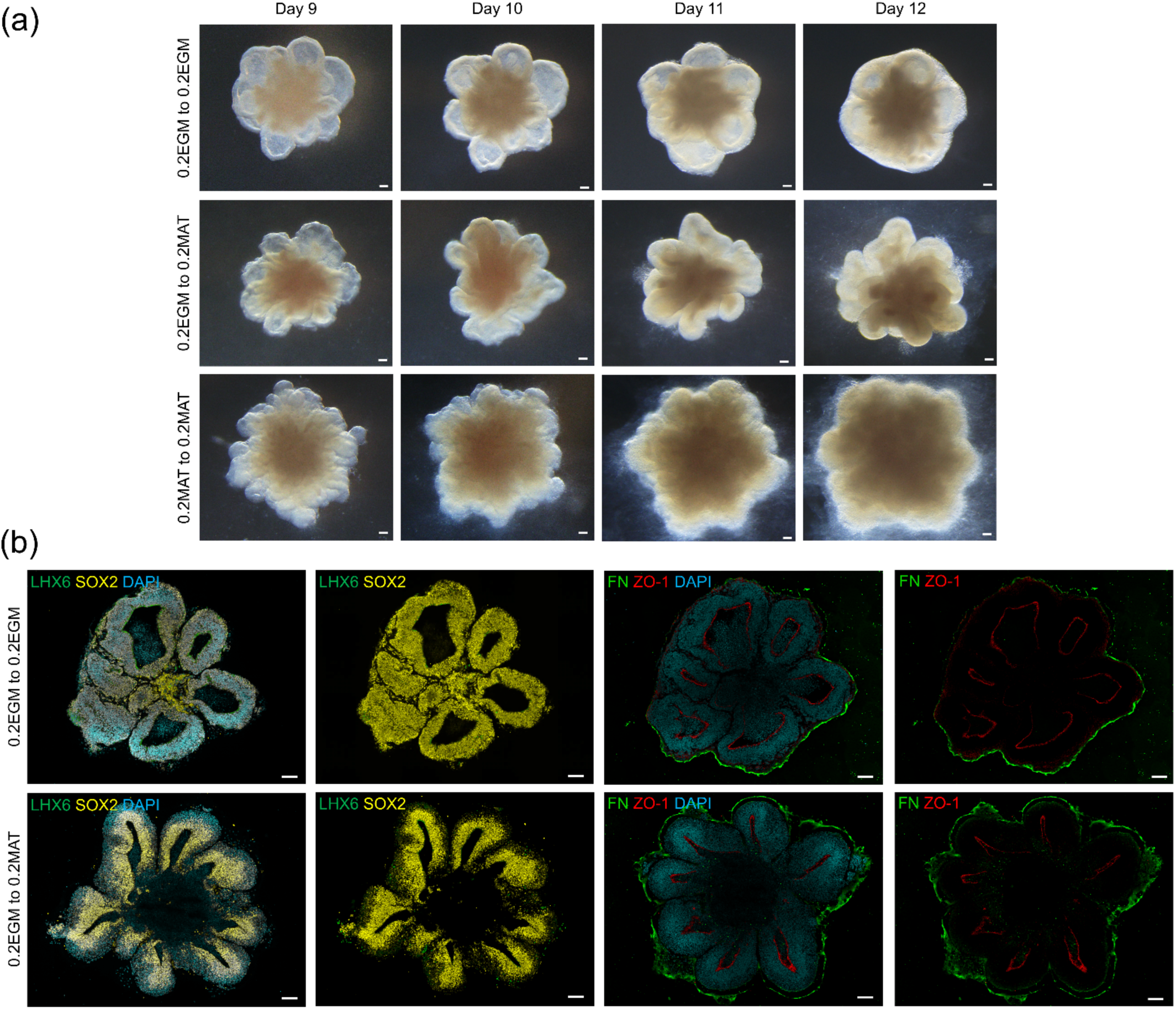
Development and characterisation of 0.2EGM D4-8 mouse ventral organoids. **(a)** Day 8-12 culture of EGM-cultured mouse ventral organoids in 0.2EGM and 0.2MAT. 0.2EGM to 0.2EGM organoids maintained thin-walled neuroepithelium. **(b)** lmmunostaining of mouse ventral forebrain organoids cultured to Day 12 in 0.2EGM revealed maintenance of SOX2+ cells throughout organoid without differentiation. Mouse ventral forebrain organoids cultured in 0.2MAT from Day 8-12 demonstrated characteristic differentiation of outer layer of neuroepithelial wall, though not to LHX6+ cell type. FN detachment from outer surface of organoid was also detected in 0.2EGM to 0.2MAT organoids but not as clear in undifferentiated 0.2EGM to 0.2EGM organoids. Scalebars: 100 µm.

**Supplementary Figure S6.**
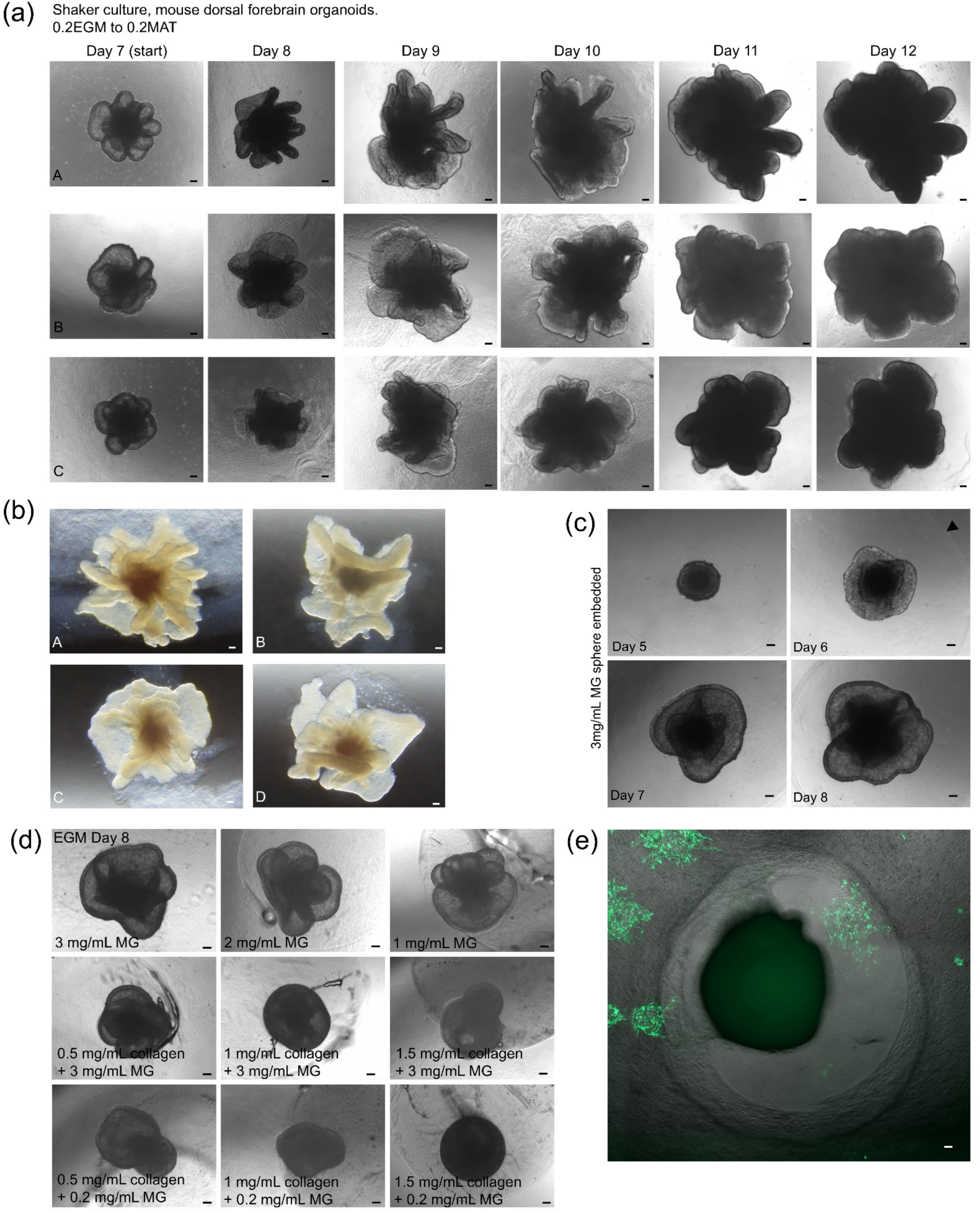
Non-coated dorsal forebrain organoids leads to a collapse ventricular lumen when cultured in Maturation medium. **(a)** Organoids cultured in 0.2EGM (Day 4-7) and moved to shaker culture in 0.2MAT (Day 7-12) display collapse of ventricular lumen and rapid proliferation. **(b)** Similar collapse of ventricular lumen was observed for 1EGM to 0.2MAT conditions (Day 9 shown). **(c)** EGM-driven expansion of MG sphere-embedded organoid. Arrowhead highlights edge of MG sphere. **(d)** Addition of collagen at 0.5, 1, and 1.5 mg/ml to 3 mg/ml MG spheres for D4-8 EGM culture had detrimental effect upon organoid bud expansion. Similar responses were observed in collagen gels containing 0.2 mg/ml MG. **(e)** EGM-expanded organoids embedded in 2 mg/ml fibrin hydrogel rapidly digest surrounding gel upon differentiation, suggesting serine protease production. Scalebars: 100 µm.

**Supplementary Figure S7.**
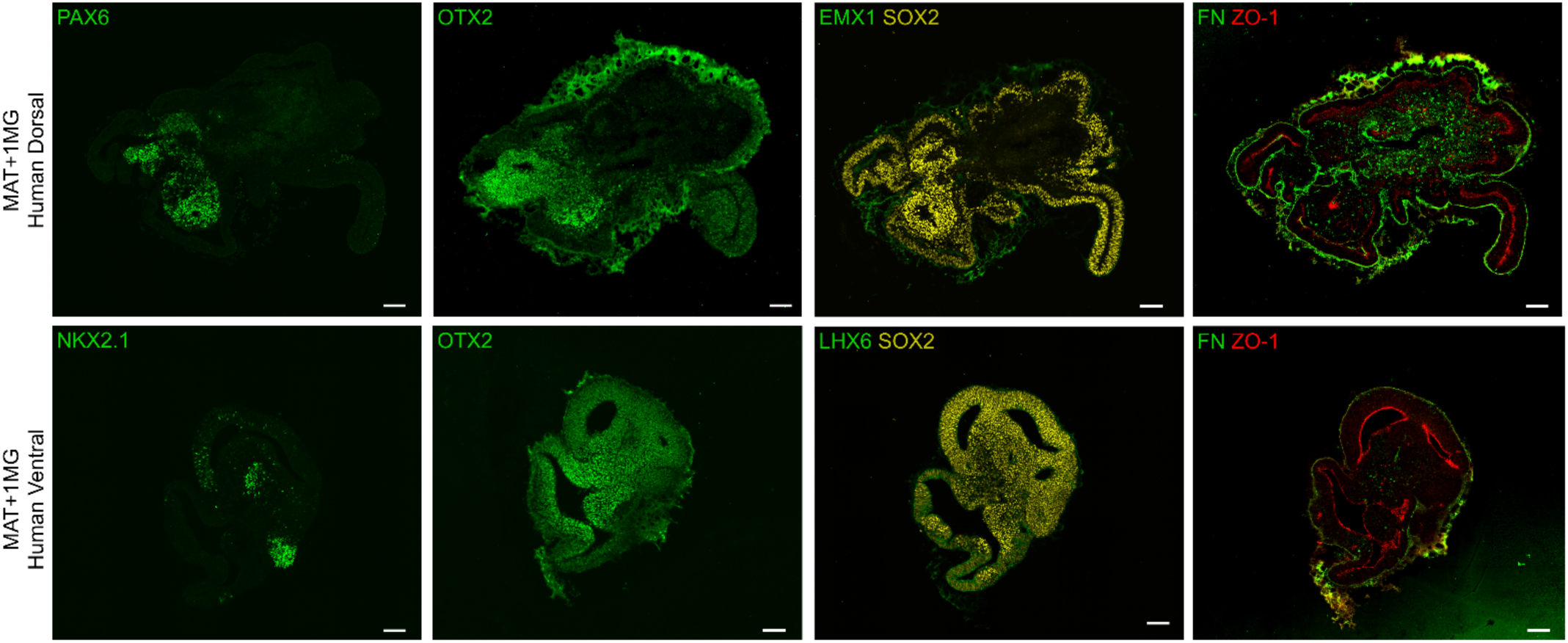
Additional human organoid data. Human organoids cultured in MAT medium from Day 7-19. Scalebars: 100 µm.

**Supplementary Figure S8.**
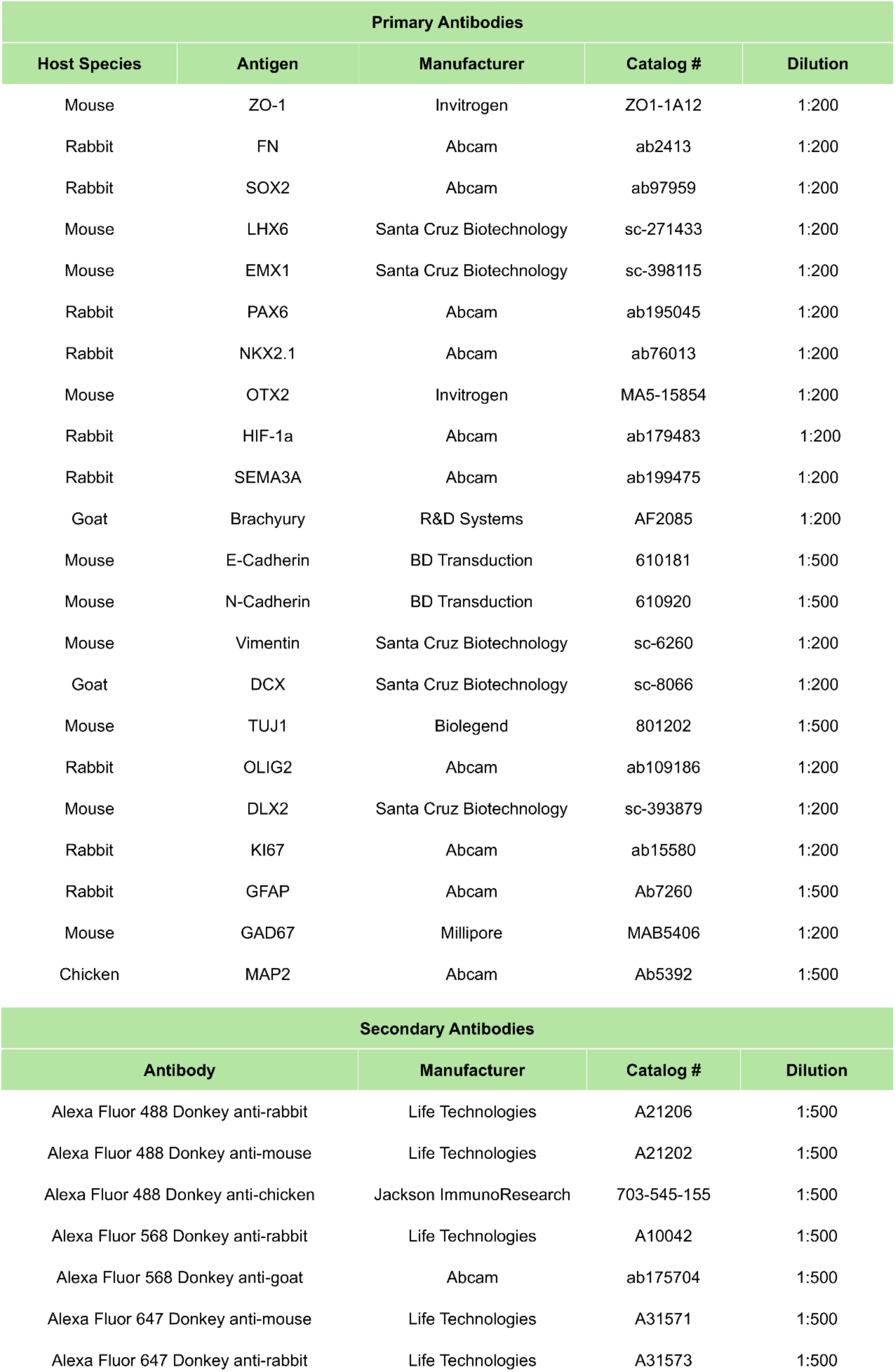
Primary and secondary antibodies.

